# Muscarinic Suppression of BK Channels in Type II Vestibular Hair Cells of Mouse Cristae

**DOI:** 10.64898/2026.04.03.716330

**Authors:** Jason M. Cote, Soroush G. Sadeghi

## Abstract

Cholinergic efferent neurons modulate sensory signaling in the peripheral vestibular system, but the cellular mechanisms underlying this modulation remain incompletely understood. In mammalian vestibular organs, type II hair cells (HC-II) receive efferent input and express α9α10-containing nicotinic acetylcholine receptors (nAChRs) that activate SK potassium channels and produce rapid hyperpolarization. Here, we examined the functional role of mAChRs in mouse vestibular HC-II using whole cell patch clamp recordings in whole tissue preparations of crista ampularis (P13–P17, male and female mice). Activation of mAChRs with oxotremorine-M inhibited voltage dependent outward currents, with the largest effects at depolarized membrane potentials. Further experiments revealed that this effect was mediated by inhibition of large conductance potassium (BK) channels: the BK antagonist iberiotoxin mimicked and occluded the muscarinic effect and muscarinic suppression was absent in mice with BK channel mutations. In contrast, blockade of SK channels with apamin did not prevent the muscarinic effect, indicating that mAChR signaling specifically targets BK mediated currents. In current clamp recordings, mAChR activation enhanced depolarization during strong current injections, consistent with increased hair cell excitability when BK channels were suppressed. These findings identify a previously unrecognized muscarinic efferent pathway in vestibular hair cells and reveal complementary cholinergic mechanisms that suppress responses to weak stimuli while enhancing responses to strong stimulation, providing a cellular basis for dynamic gain control in the vestibular periphery.

**Significance statement:** Vestibular efferent signaling shapes how head movements are encoded, but its cellular mechanisms are incompletely understood. While nicotinic acetylcholine receptors are known to reduce excitability of type II vestibular hair cells (HC-II) via small conductance (SK) channels, the role of muscarinic receptors has remained unclear. Here we show that muscarinic receptor activation selectively inhibits large conductance (BK) potassium channels in HC-II, enhancing excitability during strong depolarization. This muscarinic pathway is mechanistically distinct from nicotinic signaling and operates at a different voltage range. Together, these findings reveal a dual efferent control strategy that differentially regulates hair cell responses to slow versus fast head movements, providing new insight into how the vestibular system filters sensory input across dynamic ranges.

## Introduction

Vestibular sensory organs in the inner ear encode head motion and orientation relative to gravity, relay this information to the brainstem, and in turn receive cholinergic efferent feedback from brainstem neurons (Hilding and Wersall, 1962; Gacek and Lyon, 1974; Goldberg and Fernandez, 1980; Highstein and Baker, 1986; Perachio and Kevetter, 1989). In mammals, efferent activation increases the resting discharge of vestibular afferents (Goldberg and Fernandez, 1980; Plotnik et al., 2002; Sadeghi et al., 2009; Schneider et al., 2021) and reduces their sensitivity (Goldberg and Fernandez, 1980) with a larger effect on irregular fibers. Conversely, suppression of efferent activity lowers afferent baseline firing, again with a preferential impact on irregular afferents (Raghu et al., 2019).

The cholinergic efferent fibers innervate type II hair cells (HC-II) and afferent boutons that contact them as well as calyx terminals that cover the basolateral walls of type I hair cells (HC-I). Activation of α4β2-containing nicotinic acetylcholine receptors (nAChRs) on calyx terminals results in a rapid increase in firing rate of afferents (Holt et al., 2015). Activation of muscarinic AChRs (mAChRs) on calyces result in inhibition of KCNQ potassium channels and a change in afferent firing (Pérez et al., 2009; Kalluri et al., 2010; Ramakrishna et al., 2021) including a slow increase in firing rate of afferents (Holt et al., 2017; Lee et al., 2017; Sinha et al., 2024). In HC-II, activation of α9α10-containing nAChRs leads to a Ca²⁺ influx that activates small conductance Ca²⁺ activated potassium (SK) channels, resulting in rapid hyperpolarization (Poppi et al., 2018; Yu et al., 2020), similar to the mechanism described in cochlear hair cells (Glowatzki and Fuchs, 2000; Katz et al., 2004; Gómez-Casati et al., 2005). Whether HC-II also express functional mAChRs remains unresolved. Evidence for mAChR expression in HC-II has been reported in non-mammalian species: pigeon (Li and Correia, 2011), frog: (Derbenev et al., 2005) and single-cell RT-PCR studies in guinea pig suggest mAChR expression in mammalian HC-II (Yao et al., 2011). However, reported functional consequences of mAChR activation in HC-II differ across studies, species, and experimental preparations, with both depolarizing and hyperpolarizing effects observed (Li and Correia, 2011; Guo et al., 2012).

To directly test whether mammalian HC-II cells express mAChRs and to assess their function, we performed patch clamp recordings from HC-II in intact crista ampullaris preparations. Activation of mAChRs reduced an iberiotoxin (IBTX) sensitive outward current at positive membrane potentials, indicating inhibition of large-conductance potassium (BK) channels. In contrast, mAChR activation did not affect SK channel mediated currents in HC-II. These results were confirmed in mice with BK channel mutation (*slo^-/+^* and *slo^-/-^*), in which activation of mAChRs had no effect on HC-II outward currents. Finally, in current clamp, activation of mAChR enhanced HC-II depolarization around 0 mV compared to control conditions. Such depolarizations occur during large ciliary deflections associated with fast head movements (Rüsch and Eatock, 1996; Holt et al., 1999; Soto et al., 2002). Together, these findings suggest that cholinergic efferent input have two state-dependent effects on HC-II cells: they inhibit responses at lower membrane potentials through the α9 nAChR/SK channel complex while they enhance responses at higher membrane potentials by suppressing BK currents via mAChRs. We propose that this combination functions as a stimulus dependent gain control mechanism that shapes HC-II responses, with vestibular efferents adjusting excitability to enhance responses to fast head movements and suppress slower ones. Along with modulation of calyx terminals, this mechanism may help tune vestibular afferent dynamics and improve encoding of head motion.

## Methods

### Animals

All procedures were approved by the Institutional Animal Care and Use Committee of Johns Hopkins University, Baltimore, MD, and carried out in strict accordance with the recommendations in the Guide for the Care and Use of Laboratory Animals of the National Institute of Health. C57BL/6J mice were obtained from Jackson Labs and maintained by Johns Hopkins University School of Medicine animal facilities. Both male and female WT mice were used in the experiments. Breeding pairs of FVB/NJ mice lacking exon 1 were used for generating BK mutant animals. Exon 1 encodes both the pore-forming α-subunit and the translation initiation site of the BK channel (*Kcnma1* or *Slo1*) (Meredith et al., 2004). These animals were generously provided by Dr. Andrea Meredith (Department of Pharmacology & Physiology, University of Maryland, Baltimore, MD) through the laboratory of Dr. Machiko Shirahata (Department of Environmental Health Sciences, Bloomberg School of Public Health, Johns Hopkins University, Baltimore, MD, USA). Animals were genotyped by PCR and BK mutant (*Slo^-/-^* and *Slo^-/+^*) mice were used for experiments. All animals were used at postnatal days 13-17. Both sexes were used for WT and *Slo^-/+^* mice. For *Slo^-/-^* mice, we used only males as they are infertile and kept females for colony maintenance. *Slo^-/+^* mice were used regardless of sex for patch clamp experiments. Male and female WT littermates were also used as control.

### Electrophysiology recordings and experimental design

Dissection of the end organs and tissue preparation were carried out as previously described (Sadeghi et al. 2014, Ramakrishna et al. 2021, Ramakrishna and Sadeghi 2020, Yu et al. 2020). In brief, mice were deeply anesthetized with isoflurane and subsequently decapitated. The bony labyrinth was carefully excised and transferred to a cold extracellular solution. Under microscopic visualization, the bone overlying the ampullae of the horizontal and anterior semicircular canals, as well as the utricle, was opened. The membranous labyrinth was then isolated and dissected to expose the neuroepithelium of the two cristae and the utricle. The preparation was secured under a pin attached to a coverslip, transferred to the recording chamber, and perfused with extracellular solution at 0.5 ml/min. The extracellular solution contained (in mM): 5.8 KCl, 144 NaCl, 0.9 MgCl2, 1.3 CaCl2, 0.7 NaH2PO4, 5.6 glucose, 10 HEPES, 300 mOsm, pH 7.4 (NaOH). The intracellular solution contained (in mM): 135 KCl, 0.1 CaCl2, 5 EGTA, 5 HEPES, 2.5 Na2ATP, 290 mOsm, pH 7.2 (KOH). These solutions generate a liquid junction potential of −4 mV (Sadeghi et al., 2014) and all reported voltages were corrected accordingly in the figures and text. Drug solutions were freshly prepared on the day of the experiment from frozen stock aliquots and bath-applied using a gravity driven flow pipette (1.6 mm diameter) positioned near the tissue and connected to a VC-6 channel valve controller (Warner Instruments). Oxotremorine-M (Oxo-M; muscarinic acetylcholine receptor agonist, 20 μM), iberiotoxin (IBTX; BK channel antagonist, 150 nM), and apamin (Apa; SK channel antagonist, 300 nM) were obtained from Tocris Bioscience.

Whole cell patch clamp recordings were obtained from HC-II located in the anterior or horizontal canal cristae. Tissue was imaged using a 40× water immersion objective (NA 1.0, working distance 2.5 mm) mounted on a Zeiss Examiner D1 equipped with differential interference contrast (DIC) optics. Images were displayed on a monitor via a NC-70 video camera (Dage-MTI). Patch-clamp pipettes were fabricated from borosilicate glass capillaries (1 mm inner diameter; World Precision Instruments, 1B100F-4) using a multistep horizontal puller (Sutter P-1000) and subsequently fire polished to resistances of 7-8 MΩ. All experiments were conducted at room temperature (23–25 °C). Data were acquired using pCLAMP 10 in combination with a MultiClamp 700B amplifier (Molecular Devices), low-pass filtered at 10 kHz, and digitized at 50 kHz using a Digidata 1440A.

HC-II were visually identified by their cylindrical morphology and absence of calyx afferent terminals under DIC optics. Following membrane rupture, recordings were allowed to stabilize for 5 min with a holding potential of -74 mV to minimize time dependent changes (Ramakrishnan et al., 2021; Hurley et al., 2006). In voltage clamp, a 10 mV hyperpolarization step from −74 mV (50 ms in duration) was applied to measure membrane capacitance (Cm), membrane resistance (Rm) and series resistance (Rs). Only recordings that had Rs < 25 MΩ and a holding current of < 200 pA (average: -24.6 pA ± 5.0 pA, n = 53) were included in data analysis. Series resistance Rs was not compensated for. Resting membrane potential was recorded in current clamp mode without current injection. Identity of a recorded cell was confirmed by characteristic responses of HC-II to step depolarizations as previously described and detailed in Results section (Eatock et al., 1998; Meredith and Rennie, 2016; Poppi et al., 2018; Yu et al., 2020; Martin et al., 2024). Our voltage clamp protocol consisted of an initial hyperpolarizing step to -124 mV for 250 ms and a subsequent step for 500 ms from −124 mV to +16 mV in 20 mV increments. Baseline recordings were obtained 5 min after the beginning of intracellular recording (i.e., time 0), then pharmacological agents were applied, and data were collected 15 min after the onset of each drug application to ensure consistent effects and little variability between different cells. During current clamp recordings, the resting membrane potential was first measured with no current injection. Current injection was adjusted to hold the membrane potential between −64 mV to −74 mV for at least 10 s. The recorded HC-II was then depolarized by injecting step currents of 250 pA, 500 pA, 750 pA, and 1000pA. The steps were 2-2.5 s in duration with ∼5 s between steps so that the membrane potential returned to the initial resting value.

Time dependent changes in membrane currents during prolonged intracellular patch clamp recordings may contribute to variability, as reported previously (Hurley et al., 2006; Ramakrishna et al., 2021). To address this possibility, we performed voltage clamp control experiments in a subset of cells in which no pharmacological agents were applied throughout the recording period. Voltage step protocols were delivered at 5, 20, and 35 min after the beginning of intracellular recording (i.e., time intervals used in drug experiments) to evaluate recording stability and to confirm that any observed effects during drug application were not attributable to time dependent alterations in membrane properties.

### Data analysis

Recordings were analyzed by Clampfit 10 (Molecular Devices). For voltage steps, current amplitudes measured relative to the baseline current prior to step onset were averaged over the final 100 ms of each step. During current clamp steps, membrane voltage measured relative to the initial resting potential was averaged over the last 100 ms of the step. Results were compared before and after drug application.

### Statistical analysis

For statistical analysis, GraphPad Prism (version 11) software was used. Results are reported as mean ± SE, unless otherwise specified. Outward current responses without drug application were analyzed with one-way repeated measures ANOVA with time as the factor. Outward current responses with drug application were analyzed with two-way repeated measures ANOVA with membrane potential and treatment (baseline or drug applied) as factors. Bonferroni post hoc test was used if necessary. Either an independent samples (unpaired) or paired t-test was used for comparison between two groups. Level of statistical significance was set at α = 0.05. Validity of the assumption of normality was assessed by Shapiro-Wilk test and Quantile-Quantile (QQ) plots. Results of the Shapiro-Wilk test indicated no significant violation of normality assumption for any of the data sets analyzed.

## Results

To examine whether muscarinic acetylcholine receptor (mAChR) activation modulates membrane properties in mouse type II vestibular hair cells (HC-II), we performed whole cell patch clamp recordings in excised whole tissue preparations of the anterior and horizontal cristae from P13-17 mice. Type I HCs and type II HCs are intermingled in the crista. To preselect putative type II HCs for recordings, HCs were chosen that lack a surrounding calyx terminal. However, the final distinction between type I HCs and type II HCs was made based on their response to our voltage step protocol. In particular, type I HCs show large (in the nA range), slowly inactivating current (I_K,L_) in response to the initial hyperpolarizing voltage step from −74 mV to −124 mV due to a delayed rectifier potassium channel (Rüsch et al., 1998; Meredith and Rennie, 2016; Yu et al., 2020; Martin et al., 2024). In contrast, in type II HCs, the same hyperpolarizing step activated smaller noninactivating inward currents of a few hundred pA (Yu et al., 2020) that most likely consist of inward rectifier currents (Levin and Holt, 2012) and a hyperpolarization-activated current (*I*_h_) (Rüsch et al., 1998; Horwitz et al., 2011; Yu et al., 2020; Martin et al., 2024). Furthermore, HC-II showed characteristic voltage responses to step depolarizations (Fig. 1A) including (1) small currents in response to voltage steps near their resting membrane potential (i.e., steps between -84 mV and -64 mV) and (2) large outward currents (up to 3 nA) during depolarizing steps. On average, the resting membrane potential of type II HCs was −62 ± 10 mV (n = 18) and their membrane resistance was 541.2 ± 39.5 MΩ (n = 53), similar to the values reported previously (Eatock et al., 1998; Rüsch et al., 1998; Meredith and Rennie, 2016; Yu et al., 2020).

**Figure 1.**
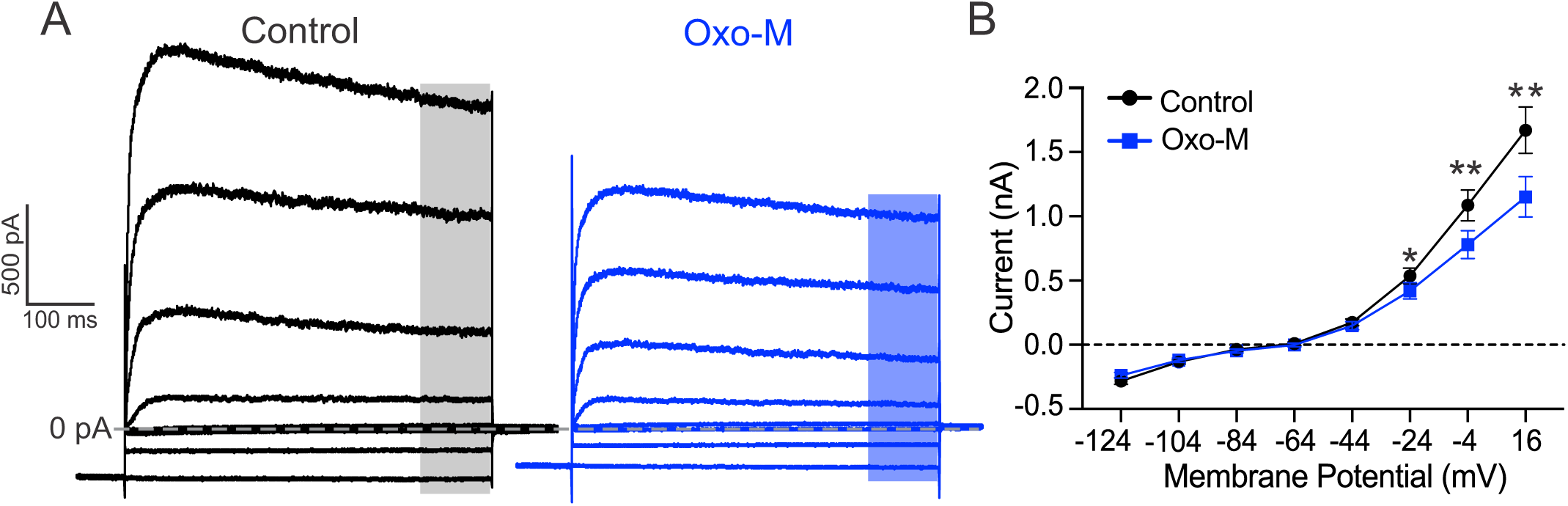
Muscarinic receptor activation suppresses voltage dependent outward currents in HC-II. **(A)** Representative outward current responses evoked by depolarizing voltage steps in a HC-II under control conditions and during application of the general muscarinic acetylcholine receptor agonist oxotremorine-M (Oxo-M; 20 μM). Cells were held at −74 mV and subjected to voltage steps from −124 mV to +16 mV in 20 mV increments. Dashed line marks 0 pA current. **(B)** For the population of recorded HC-II (n = 15) Oxo-M significantly suppressed outward currents at −24 mV, -4mV and +16 mV compared with control. * Represents significant differences at p < 0.05, and ** represents p < 0.0001

### Activation of mAChRs attenuates voltage dependent currents in HC-II

To determine whether mAChRs are functionally expressed in mouse vestibular HC-II, we applied the non-selective mAChR agonist oxotremorine methiodide (Oxo-M; 20 µM) during whole cell patch clamp recordings. Baseline control recordings were obtained 5 min after membrane rupture to ensure recording stability as previously described (Ramakrishna et al., 2021). Bath application of Oxo-M reduced voltage dependent outward currents (Fig. 1A), with large effects observed at the more depolarized membrane potentials (−24 mV, -4 mV and +16 mV). Two-way repeated-measures ANOVA (n = 15), with Membrane Potential and Treatment (Control vs. Oxo-M) as factors revealed significant main effects of Membrane Potential [F(7, 98) = 91.04, p < 0.0001] and Treatment [F(1, 14) = 18.80, p = 0.0007], as well as a significant interaction between the two factors [F(7, 98) = 25.51, p < 0.0001]. Bonferroni post hoc comparisons showed that Oxo-M significantly reduced mean outward currents at −24 mV (p = 0.029), at -4 mV (p < 0.0001), and at +16 mV (p < 0.0001) (Fig. 1B). No significant effects were observed at membrane potentials more negative than −24 mV.

These findings demonstrate that functional mAChRs are expressed in vestibular HC-II and that their activation selectively suppresses voltage sensitive outward currents at more depolarized membrane potentials. Given the activation kinetics of large-conductance Ca²⁺-activated K⁺ (BK) channels at depolarized voltages and previous reports linking muscarinic signaling to BK channel modulation in guinea pigs (Guo et al., 2012), we next investigated whether BK channels contribute to the Oxo-M induced current suppression.

### Muscarinic suppression of outward currents is mediated by inhibition of BK channels

To determine whether BK channels are suppressed by Oxo-M, we performed pharmacological occlusion experiments using the selective BK channel blocker iberiotoxin (IBTX, 150 nM). Following 5 min of stabilization post membrane rupture, a control recording was acquired to establish baseline responses. IBTX was then bath applied for 15 min to have stable results between different recordings (Ramakrishna et al., 2021), followed by 15 min of application of a cocktail containing both IBTX and Oxo-M. While IBTX application resulted in a reduction of outward current amplitudes at depolarized steps, subsequent application of IBTX and Oxo-M did not produce additional suppression (n = 8) (Fig. 2A). Two-way repeated-measures ANOVA with factors “Membrane Potential” and “Treatment” (Control, IBTX, IBTX + Oxo-M) revealed significant main effects of Membrane Potential [F (7, 49) = 66.01, p < 0.0001] and Treatment [F (2, 14) = 8.13, p = 0.0045], as well as a significant interaction [F(14, 98) = 14.67, p < 0.0001] (Fig. 2B). IBTX significantly suppressed outward currents compared to control condition at -4 mV and +16 mV (p < 0.0001, Bonferroni post hoc test). Importantly, no significant difference was observed between IBTX and IBTX + Oxo-M at any voltage step, indicating that Oxo-M does not produce additional suppression when BK channels are blocked. These findings suggest that BK channels mediate the Oxo-M sensitive component of the outward current in HC-II. The lack of additive suppression following co-application of IBTX and Oxo-M indicates that IBTX (directly) and Oxo-M (indirectly via mAChR activation) converge on a common target.

**Figure 2.**
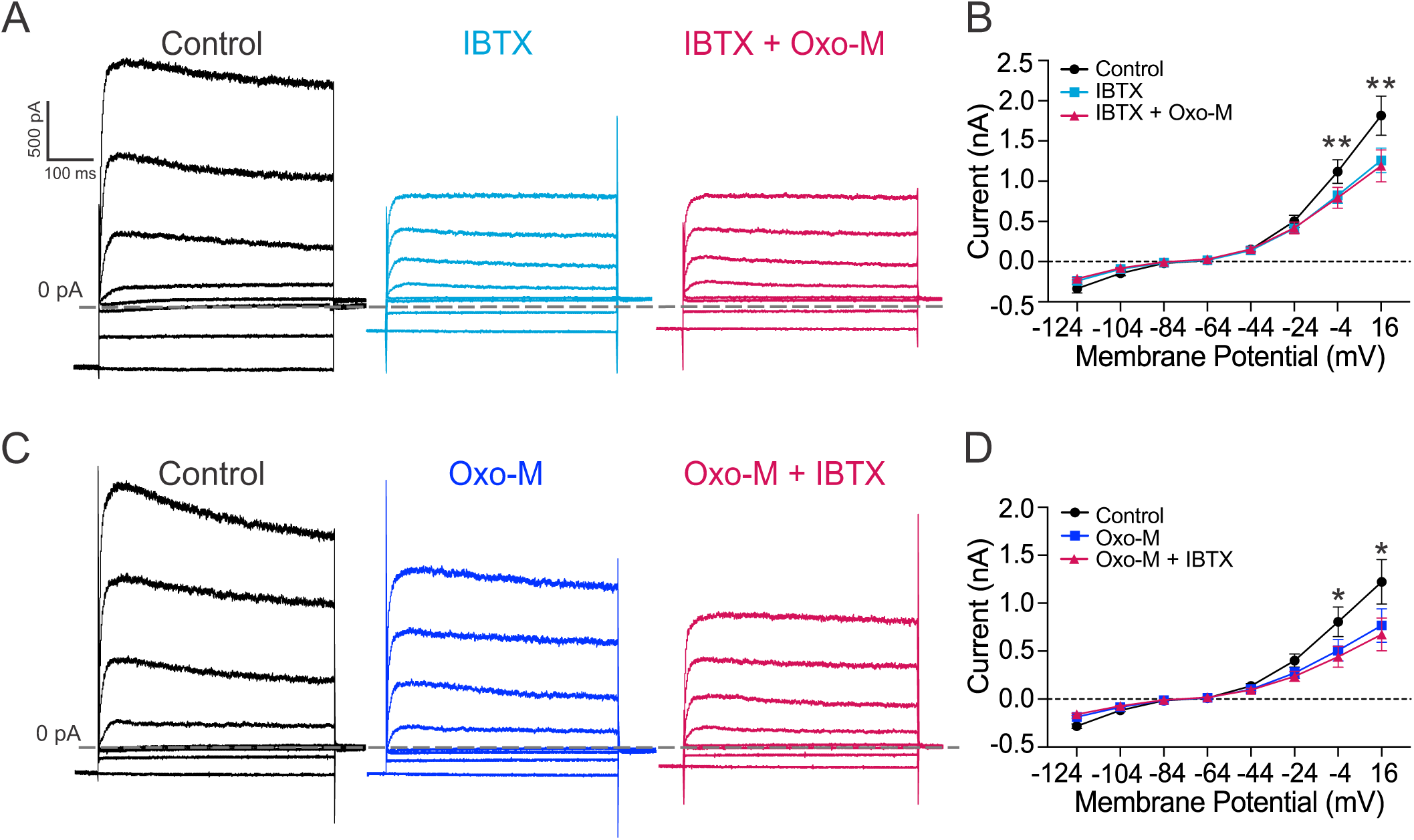
Muscarinic suppression of outward currents is mediated through inhibition of BK channels. **(A)** Representative outward current responses from a HC-II under control conditions, during application of the BK channel blocker iberiotoxin (IBTX; 150 nM), and during co-application of IBTX and Oxo-M. **(B)** For the population of recorded HC-II (n = 8), IBTX alone reduces outward currents at depolarized potentials and no additional suppression is observed with addition of Oxo-M. **(C)** Reverse occlusion experiments in which Oxo-M was applied prior to application of Oxo-M and IBTX similarly show no additional suppression upon BK channel blockade. **(D)** For the population of recorded HC-II (n = 6), while there was a reduction in current at larger depolarizing steps in the presence of Oxo-M, no additional suppression was observed when IBTX was applied in the presence of Oxo-M. * Represents significant differences at p < 0.01, and ** represents p < 0.0001.

To investigate whether BK channel blockade could suppress any residual Oxo-M sensitive currents, we performed the reverse occlusion experiments in another group of HC-II (n = 6): we applied Oxo-M first, followed by a cocktail of IBTX + Oxo-M, and used the same voltage step protocol (−124 mV to +16 mV) (Fig. 2C). Two-way ANOVA with factors “Membrane Potential” and “Treatment” revealed main effects of Membrane Potential [F (7, 42) = 35.76, p < 0.0001], Treatment [F(2, 12) = 3.94, p = 0.0482] and a significant interaction [F (14, 84) = 6.171, p < 0.0001] (Fig. 2D). As expected from our previous set of experiments (Fig. 1), Oxo-M alone significantly reduced outward currents at -4 mV and +16 mV (p < 0.01, Bonferroni post hoc test). Furthermore, there were no significant differences between Oxo-M and IBTX + Oxo-M cocktail at any voltage step.

Collectively, the results of the above forward and reverse occlusion experiments provide strong evidence that BK channels fully account for the Oxo-M sensitive outward current in vestibular type II hair cells.

### Activation of mAChR does not affect SK channel activity

Previously, we and others have shown that vestibular HC-II express SK channels that are coupled to nicotinic AChRs (Poppi et al., 2018; Yu et al., 2020). As a positive control and to further assess if Oxo-M affects SK channels, in another group of cells (n = 5) apamin (300 nM) was bath applied for 15 minutes, followed by a cocktail of Oxo-M + apamin for an additional 15 minutes and the effects were evaluated by voltage steps (Fig. 3A). A two-way repeated-measures ANOVA with factors “Membrane Potential” and “Treatment” (Control, Apamin, Apamin + Oxo-M) revealed significant main effects of Membrane Potential [F (7, 28) = 144.8, p < 0.0001] and Treatment [F (2, 8) = 21.37, p < 0.0001], as well as a significant interaction [F (14, 56) = 22.58, p < 0.0001] (Fig. 3B). As expected, application of apamin alone significantly reduced currents relative to control at -4 mV and +16 mV (p < 0.0001, Bonferroni post hoc test). However, adding Oxo-M resulted in further suppression at -24 mV, -4 mV, and +16 mV (p < 0.0001, Bonferroni post hoc test). No significant differences were found between apamin and apamin + Oxo-M relative to control at more hyperpolarized potentials.

**Figure 3.**
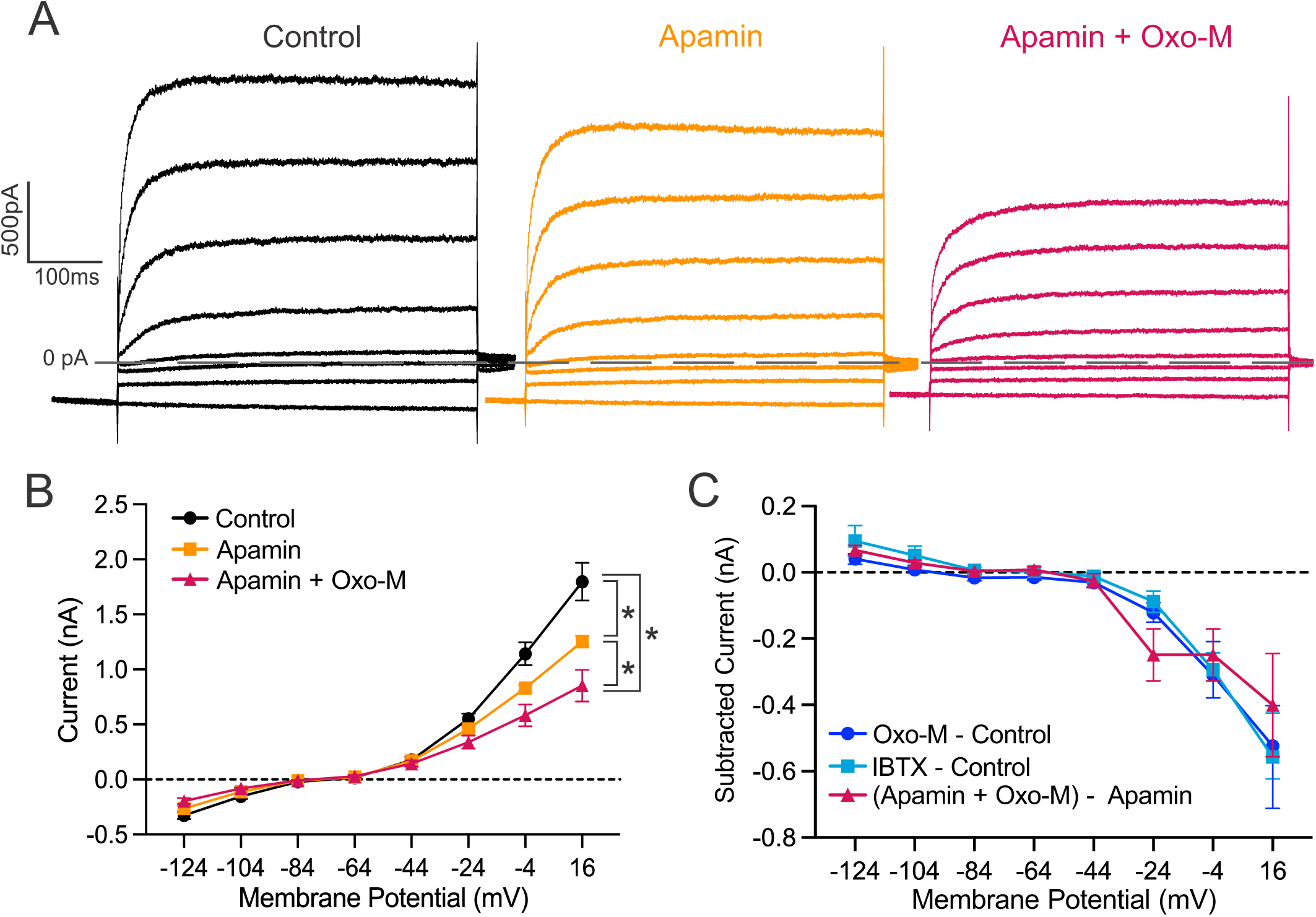
Muscarinic suppression of outward currents is not mediated through SK channels. **(A)** Representative outward current responses from a HC-II under control conditions, following application of SK channel blocker apamin (300 nM), and during co-application of apamin and Oxo-M. **(B)** For the population of recorded HC-II (n = 5), mean current–voltage relationships show that apamin reduces outward current at depolarized potentials and that additional suppression occurs with Oxo-M application. Post hoc comparisons indicate that apamin significantly reduces current relative to control at -4 mV and +16 mV (p < 0.0001) and that apamin + Oxo-M produces further suppression at -24 mV, -4 mV, and +16 mV (p < 0.0001). **(C)** Subtraction plots (drug − control) illustrate that the Oxo-M sensitive component persists in the presence of SK channel blockade, indicating that muscarinic modulation does not target SK conductances. * Represents p < 0.0001

The differences in the effects observed under the various experimental conditions (i.e., Oxo-M alone, IBTX alone, and Oxo-M following apamin) are illustrated using subtraction I–V curves (drug – control; Fig. 3C). A two-way repeated-measures ANOVA with factors “Membrane Potential” and “Treatment” revealed only a significant main effect of Membrane Potential [F (1.116, 27.91) = 36.11, p < 0.0001]. The reduction in outward current at the two most depolarized potentials was comparable across all three conditions (Bonferroni post hoc test, p > 0.05). Notably, Oxo-M continued to significantly suppress current amplitude even in the presence of apamin. This finding indicates that the Oxo-M sensitive component reflects reduced BK channel activity and is distinct from the SK mediated current.

### Activation of mAChR in HC-II has no effect in mice with BK channel mutation

To confirm that BK channels mediate the muscarinic suppression of outward currents in HC-II, we repeated the Oxo-M application protocol in BK channel mutant mice. Due to breeding constraints, most recordings were obtained from heterozygous mice (slo^+/–^, n = 7) and only two homozygous KO mice (slo^−/−^). Representative recordings (Fig. 4A) show that Oxo-M failed to alter outward currents in both the homozygous KO (Fig. 4A) and heterozygous mice (Fig. 4B). Therefore, data were pooled for group analysis (Fig. 4C). Two-way repeated-measures ANOVA with factors “Membrane Potential” and “Treatment” showed a main effect of Membrane Potential [F (7, 56) = 53.24, p < 0.0001], with no effect of treatment [F (1, 8) = 0.0005243, p = 0.9823] or an interaction [F (7, 56) = 1.617, p = 0.1497]. Thus, Oxo-M produced no significant effect at any membrane potential indicating that functional BK channels were required for the muscarinic suppression of outward currents.

**Figure 4.**
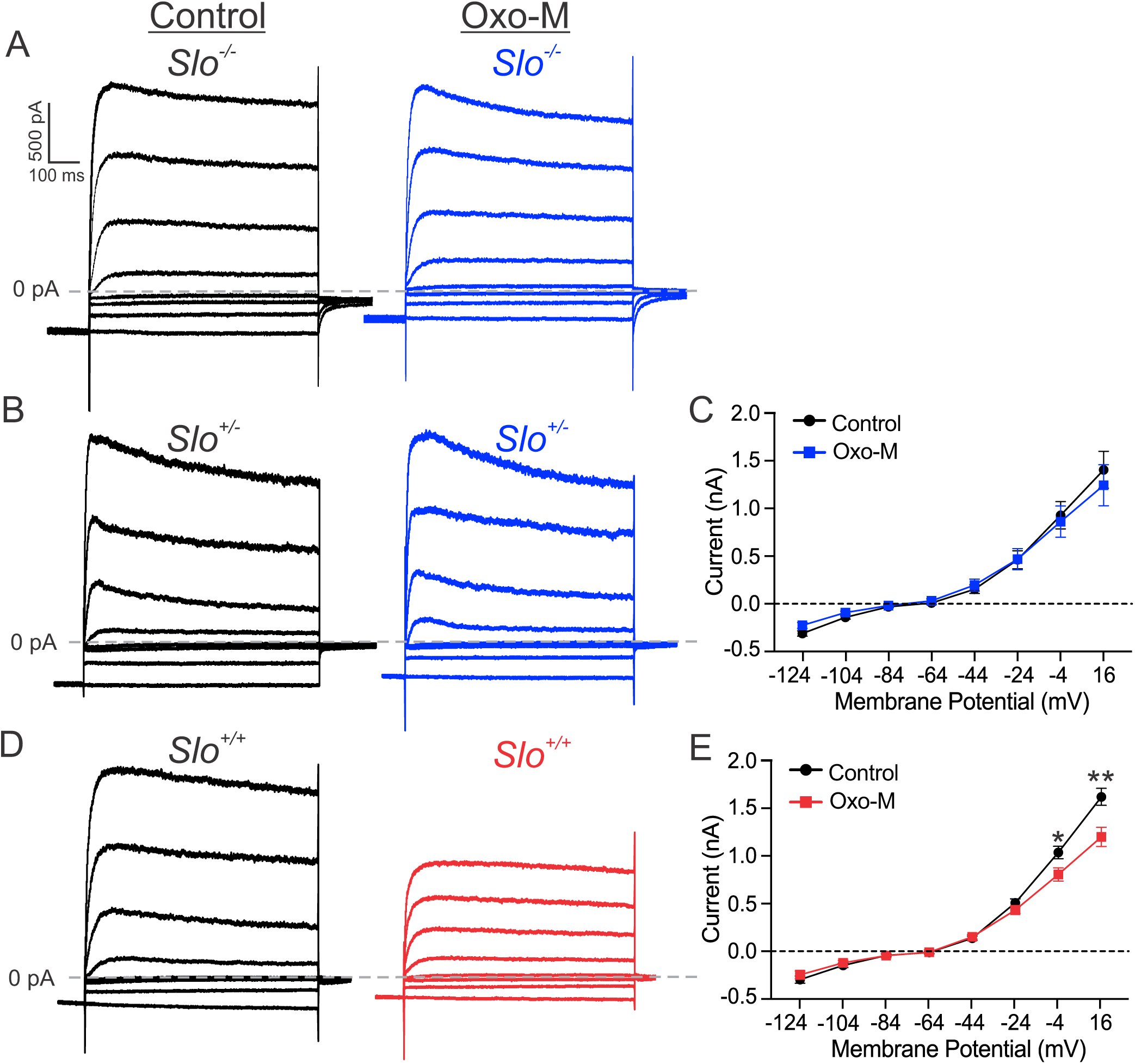
Muscarinic suppression of outward current requires BK channels. **(A)** Representative outward currents evoked by voltage step protocols in a HC-II from a BK KO (*slo^−/−^*) mouse show no effect of Oxo-M. **(B)** Representative currents from a HC-II in a heterozygous BK mutant (*slo^−/+^*) mouse similarly show no response to Oxo-M. **(C)** Pooled data from BK mutant HC-II (*slo^−/−^*, n = 2; *slo^+/−^*, n = 7) revealed no significant change in outward currents during Oxo-M application. **(D)** Representative recordings from WT littermates show robust suppression of outward currents by Oxo-M. **(E)** Population average of HC-II recordings (n = 4) from control WT littermates confirm a significant reduction in outward current at depolarized potentials following Oxo-M application. Together, these results indicate that functional BK channels are required for muscarinic suppression in HC-II. * Represents significant differences at p < 0.01 and ** represents p < 0.0001.

Since the BK mutant line was generated on a FVB background strain, which differs from the C57BL/6J background used in our pharmacological experiments, we also examined the effects of Oxo-M in WT littermates of BK mutants. As shown in a representative example (Fig. 4D), Oxo-M reduced outward currents during large depolarizing steps, similar to the effect observed in C57BL/6J mice (Fig. 1A). For the population of recorded HC-II in WT littermates (Fig. 4E, n = 4), a two-way repeated measures ANOVA with factors “Membrane Potential” and “Treatment” revealed significant effects of Membrane Potential [F (7, 21) = 678.7, p < 0.0001], Treatment [F (1, 3) = 8.986, p = 0.0578] and a significant interaction [F (7, 21) = 8.157, p < 0.0001]. Pairwise comparisons (Bonferroni post hoc test) confirmed that Oxo-M significantly reduced outward currents at –4 mV (p = 0.005) and +16 mV (p < 0.0001).

We also analyzed the baseline (‘Control’) voltage sensitive outward currents prior to Oxo-M application between BK mutants and WT littermates in a two-way repeated measures ANOVA with factors “Membrane Potential” and “Treatment”, revealing only a significant effect of Membrane Potential [F (7, 77) = 111.8, p < 0.0001]. Interestingly, there was no significant difference in baseline outward currents across voltage steps [F (1, 11) = 0.2296, p = 0.6412]. This suggests that compensatory mechanisms may preserve overall outward current amplitude in the absence of BK channels. However, the complete loss of muscarinic sensitivity in BK mutants indicates that BK channel function is specifically required for mAChR mediated modulation and cannot be functionally replaced by other potassium conductances.

Together, these results support the notion that muscarinic suppression of outward currents in vestibular HC-II is mediated by BK channels.

### Time matched controls demonstrated stable recordings

Consistent with our previous observations in calyx terminals (Ramakrishna et al., 2021), we noted that the effects of Oxo-M, IBTX, and apamin began several minutes after drug application and continued to increase over time. Therefore, all drugs were applied for 15 min in all experiments to standardize conditions and minimize variability in drug effects across recordings. To ensure that the observed effects were attributable to pharmacological manipulation rather than a side effect of the long recording times, (particularly for occlusion experiments), we performed an additional set of control experiments in C57BL6/J mice (n = 6). HC-II were recorded in voltage clamp for up to 35 min without any drug application and currents evoked by the step protocol were compared at three time points: time 0 (5 min after establishing intracellular recording) as well as 15 min and 30 min later. A two-way repeated measures ANOVA with factors “Membrane Potential” and “Treatment” across all voltage steps revealed only a significant main effect of Membrane Potential [F(7, 28) = 59.40, p < 0.0001] with no effect of treatment [F (2, 8) = 0.1431, p = 0.8688] or interaction [F (14, 56) = 1.486, p = 0.1469] (Fig 5A). Outward currents are also shown for the largest depolarizing step (Fig. 5B), which did not show a significant difference between different conditions. These results demonstrate that patch clamp recordings remained stable over time and that the observed changes in voltage sensitive currents were pharmacologically mediated.

**Figure 5.**
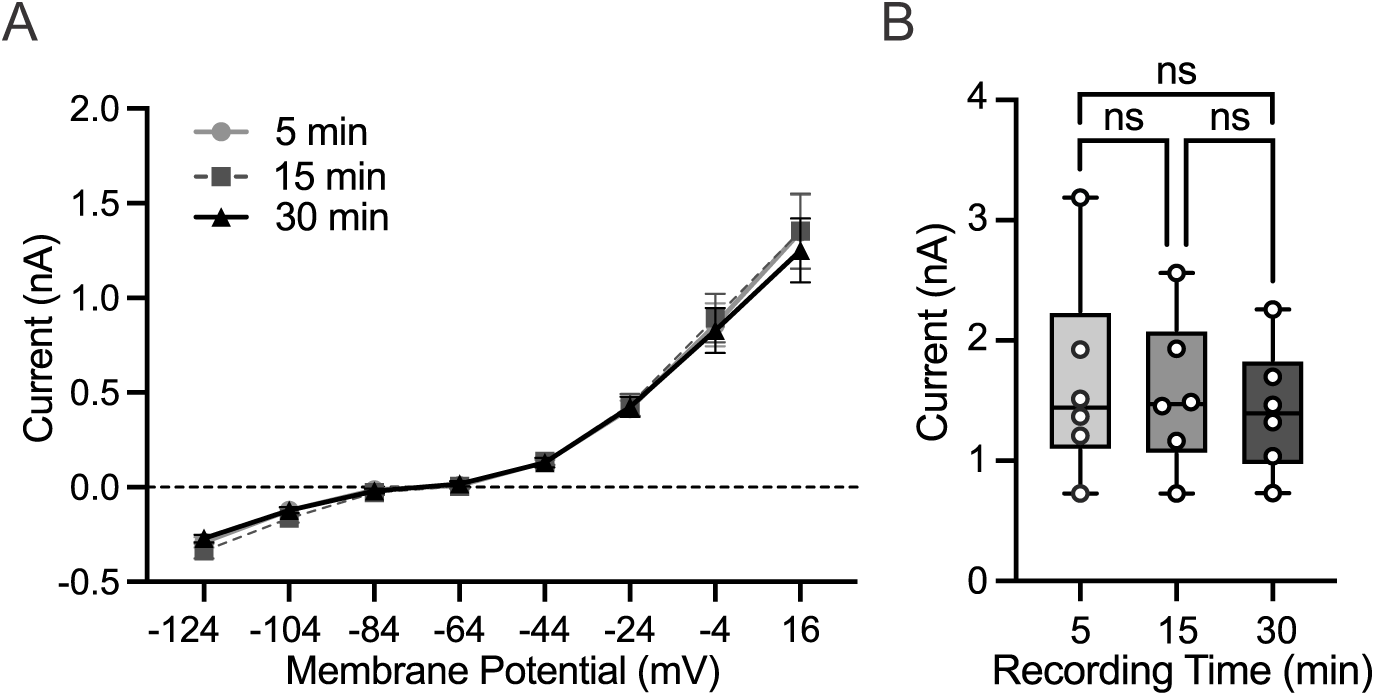
Time-matched control recordings demonstrate stability of outward currents. **(A)** Average outward currents in response to the voltage step protocol are shown for the population of recorded HC-II under control conditions at 5, 20, and 35 min after establishing whole cell configuration, in the absence of pharmacological manipulation (n = 6). **(B)** Boxplots showing comparison of outward current at the largest voltage step of 16 mV, with no significant change across time points. These results indicate that the observed drug-induced effects are not attributable to time-dependent rundown of currents. Whiskers represent 5-95% CI and box area represents the interquartile range with solid black line at median.

### mAChR Activation Enhances HC-II Depolarization

To determine whether muscarinic receptor activation alters the sensitivity of HC-II neurons to depolarizing inputs, we measured membrane potential responses to current steps ranging from 250 pA to 1 nA (250 pA increments) in current clamp mode, before and after application of Oxo-M. These injected currents served as a proxy for physiological transduction currents and were intended to depolarize HC-II to near 0 mV, the voltage at which voltage gated calcium channels mediating synaptic vesicle release exhibit maximal activity (Bao et al., 2003; Manca et al., 2021). Because some recordings exhibited a gradual depolarizing trend during the step, membrane potential was quantified as the difference in voltage during the last 100 ms of each step compared to the membrane potential before the step. Note, to have the same initial condition, current was injected to maintain the resting membrane potential near −74 mV across all recordings.

A representative example is shown in Figure 6A, where the membrane potential of a HC-II neuron reached ∼0 mV during the largest current step under control conditions. Bath application of Oxo-M increased the depolarization evoked by larger current injections compared to control. For the final 100 ms, a two-way repeated measures ANOVA with factors “Stimulation” and “Treatment” revealed significant effects of Stimulation [F (3, 15) = 30.57, p < 0.0001], Treatment [F (1, 5) = 11.34, p = 0.0199] and a significant interaction [F (3, 15) = 5.380, p = 0.0103]. Notably, pairwise comparisons (Bonferonni post hoc test) revealed significant differences only at the largest steps: 750 pA (p = 0.0078) and 1000 pA (p < 0.0001) (Fig. 6B). For the largest step, Oxo-M increased depolarization during the last 100 ms of the step by 52.79 ± 7.45 mV relative to control (Fig. 6C). Together, these findings indicate that mAChR activation enhances HC-II responsiveness specifically during strong depolarizing inputs, corresponding to larger ciliary deflections during fast head movements.

**Figure 6.**
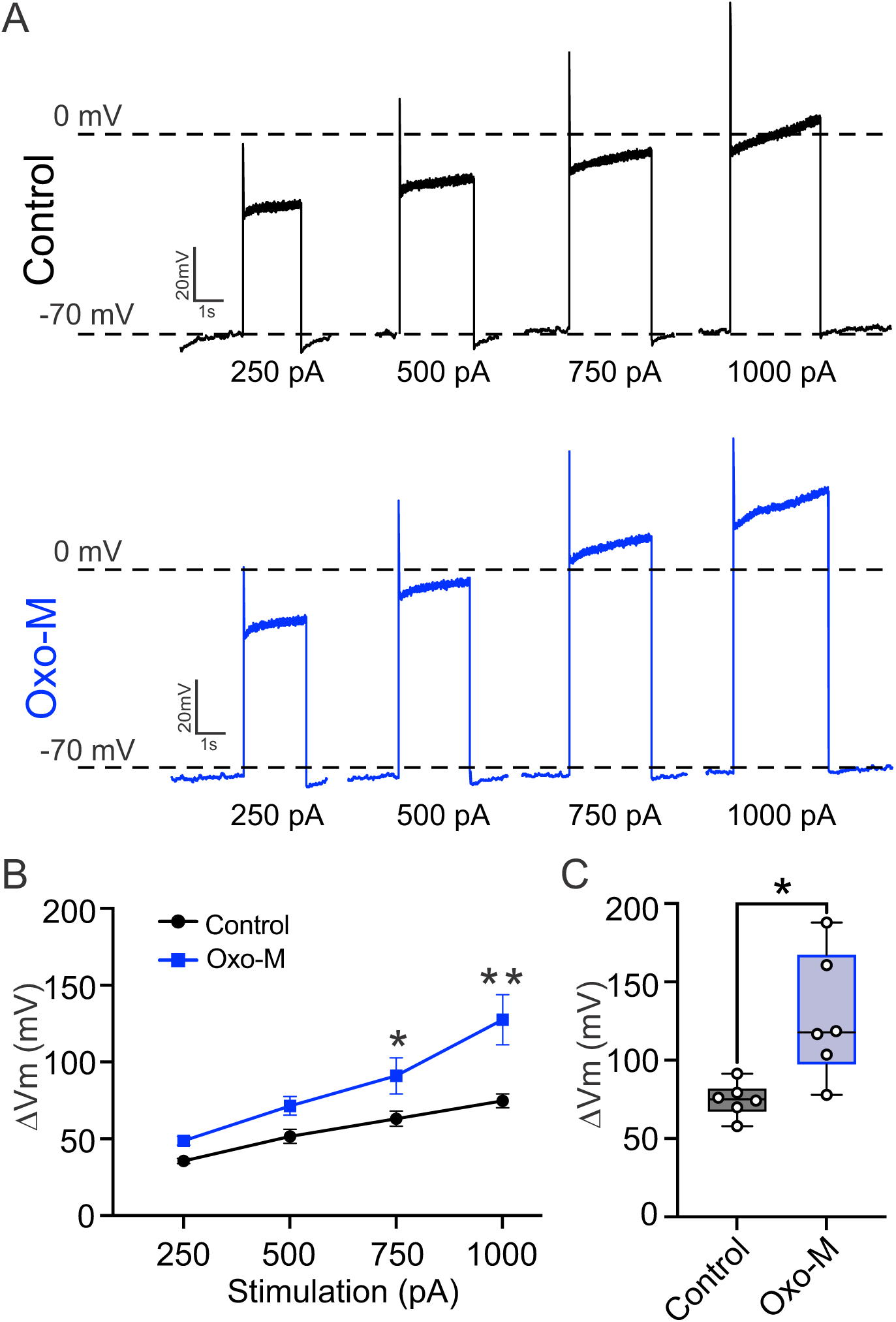
Muscarinic receptor activation enhances depolarization during current-clamp recordings. **(A)** Representative membrane potential responses from a HC-II to depolarizing current steps under control conditions and during Oxo-M application. Cells were maintained near −74 mV prior to stimulation and all data are presented as change in membrane potential (ΔmV) during the last 100 ms of the step compared to pre-step membrane potential (averaged over 100ms). Dashed lines mark 0 mV and -70 mV membrane potentials. **(B)** The population of recorded HC-II (n = 6) demonstrated significantly enhanced depolarization during Oxo-M application for the two largest current injections that drive membrane potential close to 0 mV, indicating that muscarinic activation increases HC-II excitability during strong depolarizing inputs. **(C)** Boxplots show the effect of Oxo-M on the response to the largest (1000pA) current step. Whiskers represent 5-95% CI and box area represents the interquartile range with solid black line at median. * Represents significant differences at p < 0.01 and ** represents p < 0.0001.

## Discussion

Here we show that activation of mAChRs modulates the membrane properties of mouse vestibular HC-II by suppressing BK channel conductance. Whole cell recordings from mouse cristae demonstrate that the general muscarinic agonist Oxo-M selectively reduces outward potassium currents at depolarized membrane potentials. This effect is occluded by IBTX, a BK channel antagonist iberiotoxin and is absent in BK channel knockout mice, indicating that BK channels represent the primary downstream effector of muscarinic signaling in HC-II. These results identify a previously unrecognized mechanism through which cholinergic efferents regulate the excitability of vestibular hair cells in mice.

Our findings are consistent with reports of muscarinic inhibition of voltage sensitive potassium currents in vestibular hair cells of other species, including pigeon (Li and Correia, 2011; Derbenev et al., 2005). However, they contrast with studies in guinea pig HC-II reporting that mAChR activation enhances outward currents via BK channel activation (Kong et al., 2005, 2007; Guo et al., 2012). A key methodological difference is that those studies were performed in enzymatically dissociated hair cells. Dissociation removes hair cells from the sensory epithelium and disrupts the local microenvironment, which can alter receptor signaling, ligand binding, and ion channel gating (Holt et al., 2001). In contrast, the whole tissue preparation that we used preserves the native cellular architecture and may therefore better reflect physiological signaling mechanisms.

The morphophysiological organization of potassium channels relative to efferent terminals may further explain the distinct effects of cholinergic signaling pathways in vestibular hair cells. In cochlear hair cells, α9/α10-containing nicotinic acetylcholine receptors (nAChRs) are closely co-localized with SK channels, forming a tightly coupled inhibitory complex (Glowatzki and Fuchs, 2000). Activation of these receptors permits calcium influx that rapidly activates SK channels, producing fast hyperpolarization (Glowatzki and Fuchs, 2000; Gómez-Casati et al., 2005). Similarly, vestibular HC-II express α9/α10-containing nAChRs that permit calcium entry and activate SK (but not BK) channels, producing rapid hyperpolarization at membrane potentials near –60 to –40 mV (Poppi et al., 2018; Yu et al., 2020). In contrast, the present results show that SK currents are not affected by mAChR activation, whereas BK currents are strongly suppressed. This functional segregation indicates that BK channels are located in a region separate from the nAChR–SK complex.

The differential modulation of SK and BK channels suggests that cholinergic efferents exert stimulus-dependent control over HC-II output (Fig. 7). Slow and rapid head movements produce small and large stereociliary deflections, respectively, generating correspondingly different magnitudes of mechanotransduction currents that lead to varying levels of HC-II depolarization (Eatock et al., 1998; Holt et al., 1999; Soto et al., 2002). When HC-II is modestly depolarized, SK channel activation produces rapid hyperpolarization and suppresses transmitter release. BK channels, in contrast, are activated at more depolarized membrane potentials (e.g., a fast head movement) and the inhibition of BK currents would be expected to enhance HC-II depolarization. In the present current clamp experiments, current injection served as a proxy for transduction currents, with larger injections simulating stronger bundle deflections.

**Figure 7.**
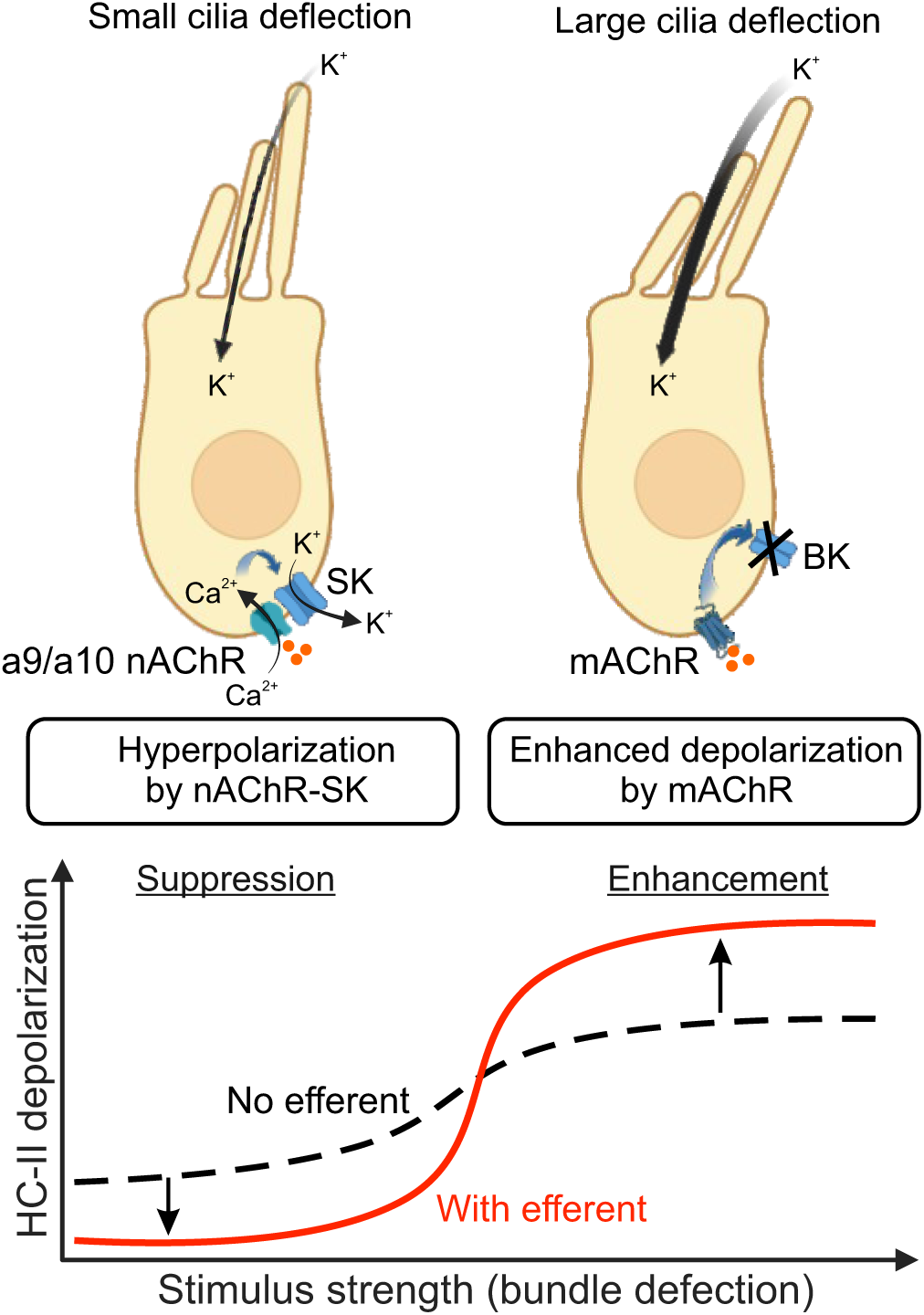
Voltage-dependent cholinergic modulation of vestibular type II hair cells. At hyperpolarized membrane potentials, such as those produced by small hair bundle deflections, activation of α9/α10 nAChRs by efferent input leads to Ca²⁺ influx and subsequent activation of SK channels. The resulting K⁺ efflux hyperpolarizes the hair cell and reduces transmitter release. At more depolarized potentials, associated with larger bundle deflections, BK channels become engaged; inhibition of BK channels via mAChRs further enhances depolarization. Together, these complementary mechanisms enable efferent input to suppress responses to weak stimuli while facilitating responses to strong stimuli, thereby modulating both the slope and the upper and lower bounds of the dynamic range, consistent with a stimulus dependent gain control of HC-II output. *Figure created in* https://BioRender.com

Consistent with BK inhibition, mAChR activation increased depolarization in current clamp recordings, particularly during larger current injections. These results suggest that muscarinic signaling selectively enhances HC-II responses during strong mechanical stimulation. The voltage dependence of these two mechanisms could therefore enable efferent input to tune HC-II output across a wide dynamic range of head movements.

Muscarinic modulation of HC-II may also act in concert with previously described effects of mAChRs on vestibular afferent terminals. In calyx terminals, mAChR activation inhibits KCNQ channels, increasing excitability and firing rate (Ramakrishna et al., 2021). Thus, muscarinic signaling may enhance responsiveness at both HC-I/calyx and HC-II/bouton synapses. Since most afferent nerve fibers receive input from both HC-I and HC-II (i.e., dimorphic terminals), mAChR activation of calyx terminals and HC-II could contribute to the increase in firing rate of afferents. However, these mechanisms may have distinct consequences for different vestibular afferent pathways. HC-II primarily communicate through quantal glutamatergic transmission, whereas type I hair cells and calyx terminals support non-quantal and possibly, quantal signaling (Songer and Eatock, 2013; Sadeghi et al., 2014; Contini et al., 2017, 2024; Spaiardi et al., 2022; Govindaraju et al., 2023; Zhou et al., 2025; Mukhopadhyay et al., 2026). Non-quantal transmission has been implicated in rapid vestibular signaling, including Vestibular sensory Evoked Potentials (VsEP) (Pastras et al., 2023; Zhou et al., 2025) and vestibulo-ocular reflex (VOR) responses (Schenberg et al., 2023; Zhou et al., 2025) In contrast, quantal transmission from HC-II contributes to sustained vestibular signals involved in gravity detection (Zhou et al., 2025). Regular afferents, which preferentially innervate the peripheral regions of the vestibular sensory epithelium containing a higher density of HC-II, are thought to encode gravitational signals (Jamali et al., 2019) and to drive VOR responses (Minor and Goldberg, 1991). By contrast, irregular afferents that innervate the central zones of the sensory epithelium contact more HC-I and encode rapid head movements (Ono et al., 2020). It is therefore conceivable that muscarinic enhancement of HC-II excitability may preferentially influence regular afferent pathways, potentially improving their responsiveness to faster head movements.

In conclusion, our results support a model in which efferent input act as a stimulus dependent gain control system that differentially shapes HC-II responses across the spectrum of head movements. In this framework, vestibular efferents do not simply inhibit peripheral activity but instead dynamically reconfigure HC-II excitability to optimize encoding of behaviorally relevant motion signals.

## Acknowledgements

This work was supported by National Institute on Deafness and Other Communication Disorders grant R01 DC019380 to SGS and R01 DC012957 to EG, T32 Postdoctoral Training Grant 90094636 to JC, David M Rubenstein Professorship to EG, David M Rubenstein Precision Medicine Center of Excellence in Hearing Fund, and Lloyd B. Minor Center for Vestibular and Skull Base Sciences.

